# Single Cells Signal Non-self from Self via ‘XOR’ and ‘NOT EQUALS’ Logic in a Dimer of Dimers

**DOI:** 10.1101/2024.10.01.616003

**Authors:** Rocky An, Manu Prakash

## Abstract

The ciliate *Tetrahymena thermophila*’s seven mating types are defined by unique receptor-ligand pairs (Mta/Mtb). While Mta and Mtb are known to participate in a mating signal complex, how they distinguish between oneself and six non-self cell types remains unknown. AlphaFold3 predictions reveal Mta/Mtb as large glycoproteins likely derived from ancient, unisexual, intercellular adhesion molecules. Since homologous binary-type systems perform XOR by switching mtA expression, we show spectrum (*n*) types naturally extend XOR to multi-bit NOT EQUALS operations via differential affinities of Mta/Mtb dimers. We model kinetics begetting the ‘*n* + 1th’ type, demonstrating self-inhibition by *trans*-homophilic Mtb-Mtb. A computational approach reconciles recent and classical evidence for mating exclusivity, including selfing failures (same type mating). Binding kinetics enables fast, robust intercellular computation across an intermembrane mating space. Thus, Mta/Mtb families are a model system allowing us to derive a ‘calculus’ of antigenic variation and inspire synthetic designs of XOR logic underlying self/non-self recognition.

**Graphical Abstract:** 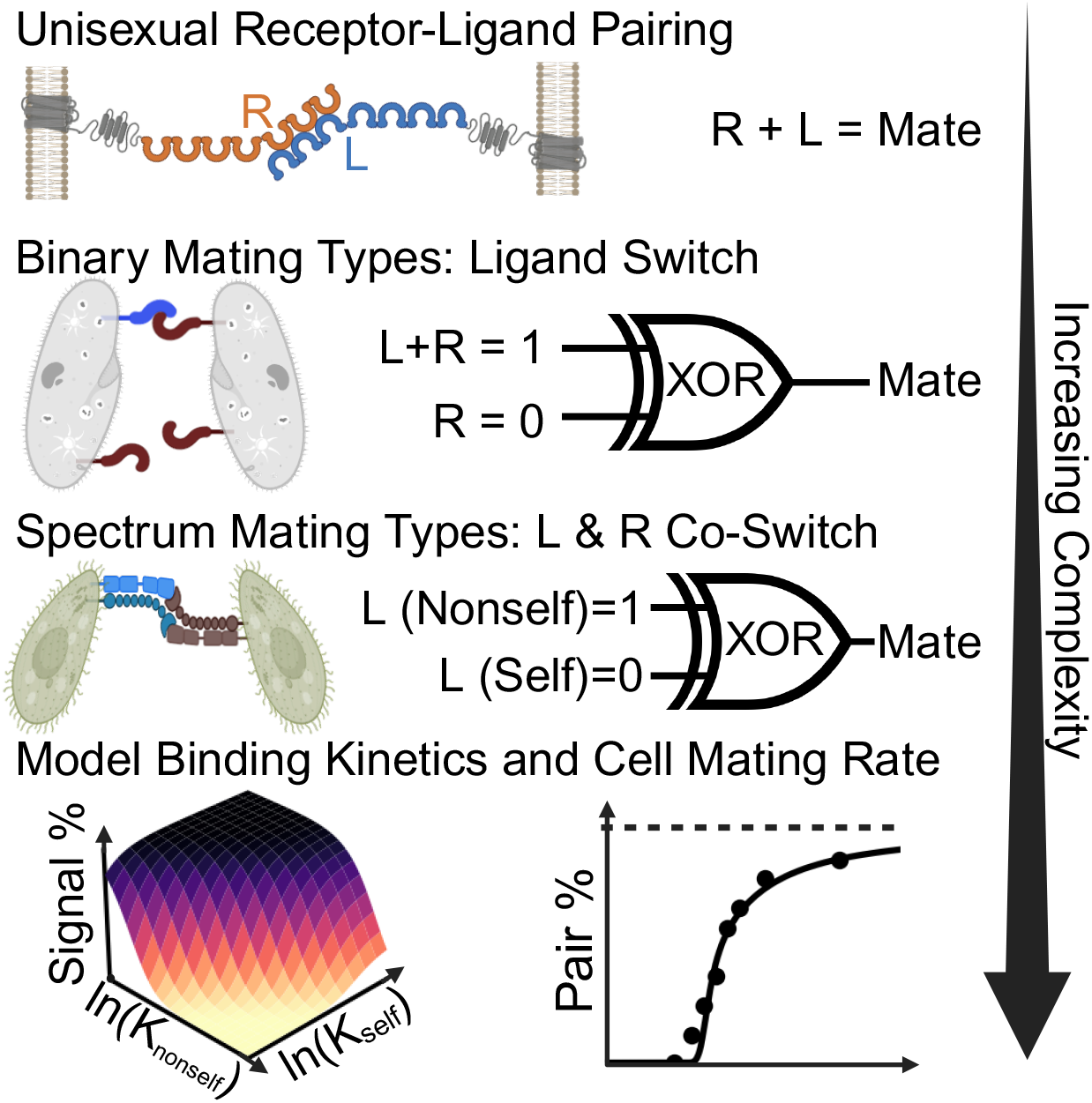

**Highlights:**

- Within minutes, swimming cells signal compatible mating types via membrane proteins
- AlphaFold insights reveal Mta/Mtb family as elongate adhesion receptors/ligands
- Relative affinities of two dimers explain multi-bit intercellular signaling capacity
- Dimer-of-dimers compute NOT EQUALS via homo-vs. hetero-philic interactions
- **In Brief** Free-living ciliates *T. thermophila* breed only when two cells are different in mating type, out of seven types total. Their mating-type-specific coexpression of a receptor-ligand glycoprotein pair reveals how competitive homophilic vs. heterophilic binding computes self vs. non-self recognition with single-layer intercellular logic. Proteins perform binary XOR gates (with two mating types) that extend to multi-bit ‘NOT EQUALS’ (with multiple mating types). The work illuminates principles of eXclusivity in molecular recognition and inspires the design of synthetic cellular recognition systems.

## INTRODUCTION

Membrane receptors can bind multiple ligands as inputs, mediating a form of intercellular signaling with the advantage of fast, single-layer computational efficiency akin to digital logic gates. By bypassing limitations typical of soluble and transcriptional systems such as concentration-dependence [1], signal delay, and oscillatory feedback [2], membrane contact-mediated signaling naturally underlies a range of computer-like complexities across neural, sexual, and immunological synapses. Many gaps in our knowledge of receptor-ligand signaling capability are about its computational limits [3], important for the logic-inspired design of engineered receptors that currently include ‘AND’, ‘OR’, and ‘NOT’ [4, 5]. How are the principles of molecular recognition built from simpler protein logic gates?

Previous research has linked receptor-ligand networks, integration of multiple signaling pathways, and environmental context to the diversity of multicellular cell types and identities [6, 7]. From a logical viewpoint, cells create combinatorial networks of AND or OR gates by exploiting gene duplication and an intrinsic promiscuity of receptors. However, some cell types seem to operate not with promiscuity, but by other principles. For example, neuronal self-repulsion is poorly understood, although known to involve a combinatorial expression of sixty protocadherin variants that *trans*-homodimerize to form a zipper-like cluster of variants [8, 9]. Knockout abolishes self-repulsion [10], implying protocadherin functions akin to a NAND gate.

The protein ‘XOR’ (eXclusive OR) gate, which necessarily requires two ligand inputs that exhibit negative cooperativity [11, 12], is more elusive. The rarity of XOR in proteins parallels the stagnation in the AI field (1970’s first AI winter) caused by the perceptron’s inability to learn the XOR function. In a similar vein, chimeric antigen receptor (CAR) T cell therapies have seen only incremental improvements with the use of ‘AND’ gate receptors [13, 14] to recognize non-self cancer antigens, facing challenges of both antigenic escape (causing negative signaling) and localized targeting of non-healthy cells (requiring positive signaling). We propose that membrane proteins can address these two competing challenges by computing XOR logic—and even higher functions—for simultaneous self plus non-self recognition. The mating proteins of the ciliates, a protist group, are examples which enable a cell to breed specifically with a partner that is different in mating type (MT). Two requirements are necessary for MT proteins: receptor-ligand signaling must be species-specific while signal inhibition must be specific against cells of the same MT.

Here, we demonstrate how two apposing cells in contact realize the XOR gate with two ligand inputs that exhibit *trans*-homomeric affinity to inhibit each other. As found in sexual *Paramecium*, this represents the simplest possible mechanism for ‘male’ and ‘female’ membrane receptors to compute XOR. In comparison, *Tetrahymena* (Box S1) encodes multiple, non-binary MT by the co-expression of a MT-specific pair of membrane proteins (Mta/Mtb). To our knowledge, no other mating system employs two membrane proteins, thus raising several questions about their mechanistic purpose. Why—in a related ciliate genus—*Paramecium aurelia* species can only encode binary MT by switching surface expression of a single ligand, named mtA, but a *Tetrahymena* Mta/Mtb pair of receptor-ligand pairs increases the number of possible MT from two to many, is a complex systems-level puzzle. In this study, we describe how XOR-driven receptors can evolve into an expanded network of MT proteins that barcode seven cell type identities that recognize self-ligands to inhibit self-mating, an overall process we define to be a ‘NOT EQUAL’ (≠) computation as the mechanistic basis for self vs. nonself determination.

### Box S1

The Unicellular Protists *Tetrahymena* Encode Multiple Mating Type Identitiesg Type Identities

The model organism *Tetrahymena thermophila* is a unicellular, free-living ciliate with rich scientific history that continues to the present day. In 1923, *Tetrahymena* was first grown in axenic culture by Lwoff, who later became a Nobel laureate alongside Jacob and Monod (who were both pioneers in the study of enzyme kinetics, regulatory signal feedback, and the *lac* operon). *T. thermophila* has served as an educational model organism [15] and been implemented for the functional expression of parasite antigens [16] as well as mammalian voltage-gated ion channels [17]. Despite shared eukaryotic commonalities, a unique aspect of *Tetrahymena* is its encoding of *clonally invariant* surface proteins that define non-binary mating types (MT)—representing a form of clonal cellular identity. All *T. thermophila* isolates were found to convey one of seven mutually exclusive MT (sometimes affectionately dubbed ‘sexes’), yet how such a prismatic spectrum of MT evolved remains a mystery. MT choice occurs once after a sexual maturation phase and is randomly influenced by environmental factors (Orias et al. [18], review the epigenetics of this process). Subsequent sequencing revealed MT to be encoded by the surface expression of two large (∼200 kDa) transmembrane proteins, Mta and Mtb, which are MT-specific due to the elimination of all 6 other MT gene pairs from the transcriptionally active macronucleus of a cell [19].

The genus *Tetrahymena* are generally pond-dwellers notorious for their adaptability to a global range of environments. Their diversity is driven by a preference to breed more opportunistically, avoid inbreeding, and outcross more frequently. This is likely enabled by multiple MT [20] compared to binary MT in *Paramecium*. For further comparison, *Paramecium* and *Tetrahymena* MT proteins have sequence homology [21]. The two genera diverged an estimated one billion years ago, but the existential and conspecific requirements of eukaryote sexual reproduction is conducive to sex-related genes being nearly as conserved as those for the cell cycle. Due to the high fidelity and tractability of unicellular sexual signaling, it is attractive to experimentally exploit their simplicity [22]. Two cells mate and fertilize each other under starvation pressure, which is replicated experimentally by placing cells in buffered solution for more than a few hours.

Non-self recognition may also manifest by different strategies apart from mating. The asexual species *T. vorax* recognizes and phagocytizes live interspecies ciliates such as *T. thermophila* [23]. Species cannibalism and self-cannibalism can be inhibited by biophysical and biochemical signals [24], but MT membrane proteins are unique by dually recognizing species-specific non-selves as well as MT-self, via physical contacts. By bridging intercellular swimming contact and behavior, MT proteins’ exclusivity and bidirectional function likely drives sexual isolation and speciation themselves [25]. Contact recognition by adhesion receptors are thought to have driven the evolution of multicellular animal adhesion [26] and possibly of cell-cell fusion [27]. Again, these are cases of only self-recognition. Various ciliates, with the same “toolbox” of organelle and protein functions [28, 29], accomplish not only binary and spectrum mating, but also a diversity of intercellular behaviors.

During sexual conjugation (mating of ciliates), *Tetrahymena* undergo a tightly coordinated process (Figure 7A) involving rapid swimming and intercellular collision-mediated signaling. Cells deciliate at the anterior adhesion junction where there is also a morphological tip-transformation analogous to schmooing in mating yeast. Although physical micromanipulation is possible to assay ciliate phagocytosis, *Tetrahymena* are relatively rigid and necessarily require ciliary swimming motility to align adhesion junctions [30, 31]. We are developing the capability of live, long-term tracking of motile single cells including *Tetrahymena* [32], which could provide future visual confirmation of our collision model (Figure 6).

We dedicate the preliminary part of our results to describe the binary MT of *P. aurelia* as a starting point to compare with the multiple MT of *T. thermophila* (Figure 1D). Contact-mediated communication necessarily involves fast and condensed signals, thus we predict ciliate protein mechanisms as computations constrained to the intermembrane space between two mating cells. While binary MT signaling can be abstracted to binary XOR of ligand inputs, multiple MT as multi-bit logic is more complicated. We model the interaction distinguishing non-self from self with a dimer-of-dimers receptor activated only by *trans*-heterophillic vs homophilic affinities. *T. thermophila* has a well-defined—yet underexplored—set of seven MT receptor/ligand pairs which provides a model for protein interactions as ‘digits’ to encode seven different MT. A combined classical and computational approach, guided by all-atom protein modeling by Google Deepmind’s AlphaFold3 [34], also informs our understanding of the variational expansion of antigens such as those encoded by pathogens. We discuss a first principles generalization of molecular recognition as a circuit of multiple bits. With a kinetic and structural lens on previous empirical work on *Tetrahymena*, we propose testable predictions that would agree with an evolution- and information-minimal mechanistic characterization of a multiple MT system. Its mechanism is robust for sex, yet minimizes the informational complexity of self/non-self determination.

**Figure 1:**
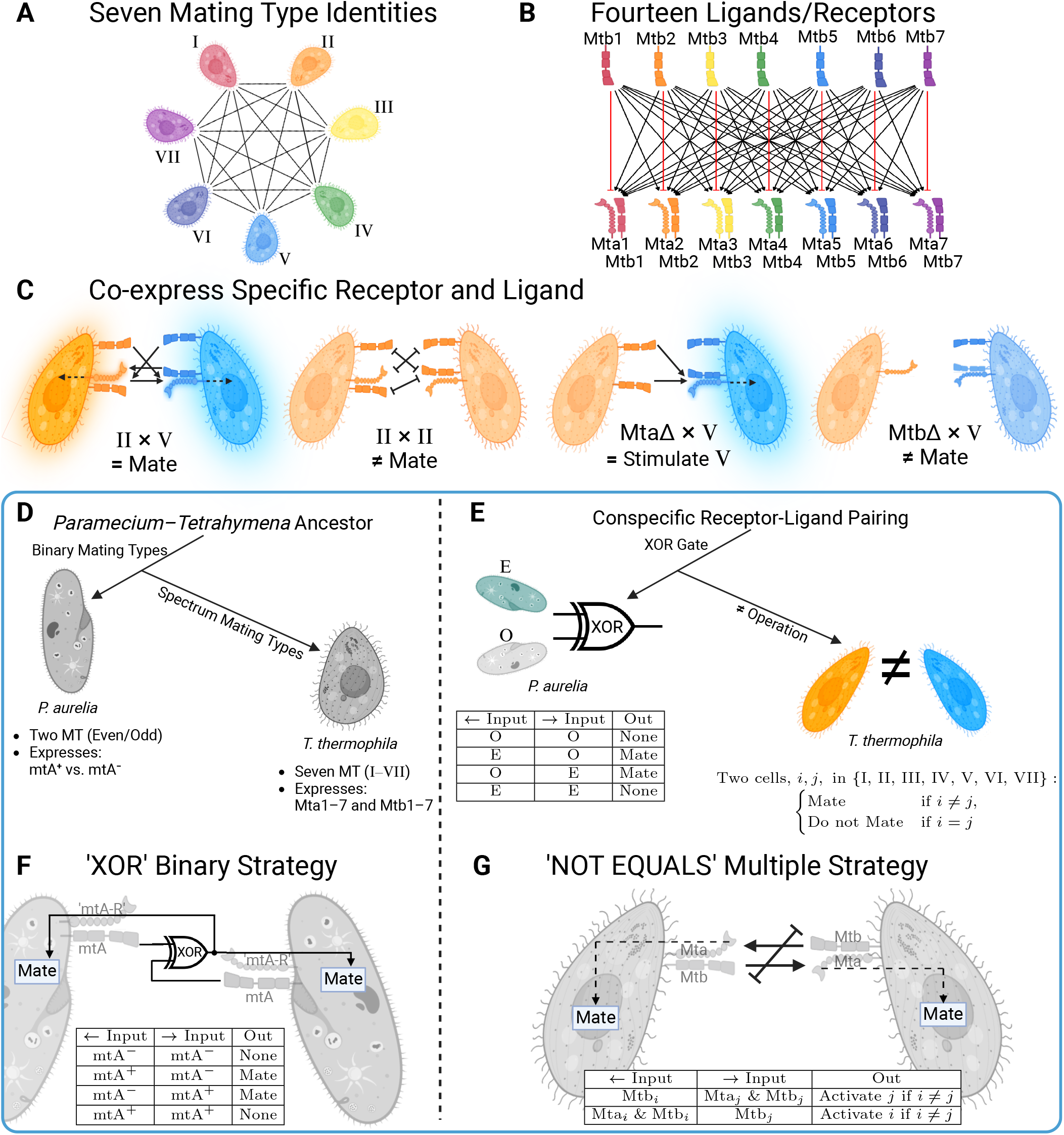
Mating type compatibility is a computation. A) Mating compatibility network. A *T. thermophila* cell line expresses one of seven mating types (MT) (colored in a spectrum) and can mate with all other MT expect its own. B) Directional representation of non-self activation by ligand (above) to receptor (below) signaling with Mta and Mtb varying per MT. C) Asymmetric stimulation in Mta knockout experiments. (MtaΔ × WT) Mta knockout cells are not stimulable, but they still activate the expression of adhesion proteins on an apposing cell (right, glowing cell). Asymmetric cells form weak pairs but both cells do not fully costimulate as they do not exhibit tip-transformation [19, 33]. For neither mixture of “mtbΔ × WT” nor “mtaΔmtbΔ × WT”, cells cannot pair nor stimulate [19]. D) The ciliate ancestor of *Paramecium* and *Tetrahymena* expressed either binary MT or was unisexual. Only the surface expression of mtA is MT-specific in the *P. aurelia* species complex. In *Tetrahymena*, both Mta coupled with Mtb are MT-specific. E) Binary MT, Even (E) and Odd (O), behavior is a XOR gate. The logic underlying two non-binary inputs we define as a NOT EQUALS (≠) operation. F) XOR membrane protein logic is based on input from each cell (left and right) via ‘ON’ vs. ‘OFF’ surface expression of mtA (mtA^+^ vs. mtA^−^). Ligand-receptor signaling catalyzes strong pairing and a subsequent mating reaction (arrow) between two cells. G) *Tetrahymena* mutual costimulation requires bilateral non-self activation (left and right arrows) and absence of self-inhibition (overlayed inhibition arrow). Mtb on one cell can activate a complementary Mta/Mtb complex on an apposing cell, causing downstream stimulation (dashed arrow).

## RESULTS

### Mating Type Signaling Shows Directionality and Mta/Mtb are Nonredundant

Mating-type recognition occurs when two swimming cells contact each other via the interaction of their mating-type receptors (Box S1). Cells form pairs after 30-60 minutes of mixing two *T. thermophila* cell lines, thus begetting a combinatorial pairing recognition network of *n*(*n* – 1)/2 = 21 mutually exclusive interactions for *n* = 7 number of MT (Figure 1A). With only two MT-specific proteins, it is difficult to realistically propose 21 unique conformational couplings. Instead, a mating receptor only needs to distinguish self from non-self and activate upon non-self ligands (Figure 1B). Theoretically, a engineered receptor could be tuned for aversion of a self-ligand plus promiscuity in six non-self interactions, but pairwise receptor-ligand affinity, alone, cannot explain several observations about the process [19, 35, 33].

Mta recognizes Mtb across the intermembrane space [33] (i.e. heterotypically *in-trans*), yet knockout studies reveal Mta and Mtb have non-redundant roles (Figure 1C). Mtb knockout renders cells unable to stimulate others or be stimulated, while Mta knockout only prevents their internal stimulation [19]. Classical work (reviewed by Cole [36]) found that fixed, dead cells can still activate MT-complementary live cells, an observation which may preclude active biochemical signaling mechanisms and further support an externally presented Mtb membrane protein as a MT identifier. It follows that Mtb presents a MT signal and an Mta/Mtb complex receives the signal on an apposing side, so we define Mtb to be an *upstream* ligand and Mta to be a *downstream* receptor. This naming scheme will be important to understand the circuit and evolutionary relationship to XOR and mating *Paramecium*.

### Adhesion Receptors Compute XOR Across Two Pairing Cells

We first consider a simplified base case where a cell must only distinguish between two MT, as observed in the *Paramecium aurelia* species complex (Figure 1D–E). A *P. aurelia* species has only binary MT (Even vs. Odd) and one MT receptor (named mtA) which is homologous to either Mta or Mtb in *Tetrahymena* [37]. The two MT function with a binary switch: E-type cells express mtA while O-type cells do not. In the well-studied *P. tetraurelia*, O-type cells excise the mtA promoter [38]. This single-protein encoding of E and O is remarkably simple but fundamentally asymmetric in mtA^+^ vs. mtA^−^ surface expression. Such asymmetric dependence of mating by the strength of interacting receptor-ligand pairs is generally believed to lead to evolutionary stability only toward binary-types [25]. This asymmetry archetype, likely inspired by sperm-egg interactions, is thought to pressure single-cell signaling towards a binary MT system. However, we demonstrate that a symmetric requirement of MT protein expression can give rise to multiple MT.

To investigate the structural grounds of binary MT determination in *Paramecium tetraurelia*, we employed AlphaFold3 [34] to predict structures (Figure 2A) of mtA as well as the three homologs named mtA-Like (mtAL1-3) that are induced by starvation [37]. The mtA family consists of five transmembrane helices, a cysteine-rich, growth factor receptor-like (GFR-like) domain, and seven extracellular β-sandwich domains. Their extracellular domains are immunoglobulin (Ig)-like and form an elongate structure, thus sharing characteristics of cellular adhesion molecules quintessential for intercellular contact recognition. Notably across *P. aurelia* species, mtA and mtAL2 also feature hydrophillic cytoplasmic tails which may have signaling or localization functions, as mtA contains a known C-terminal ‘VxPx’ ciliary targeting motif [39].

**Figure 2:**
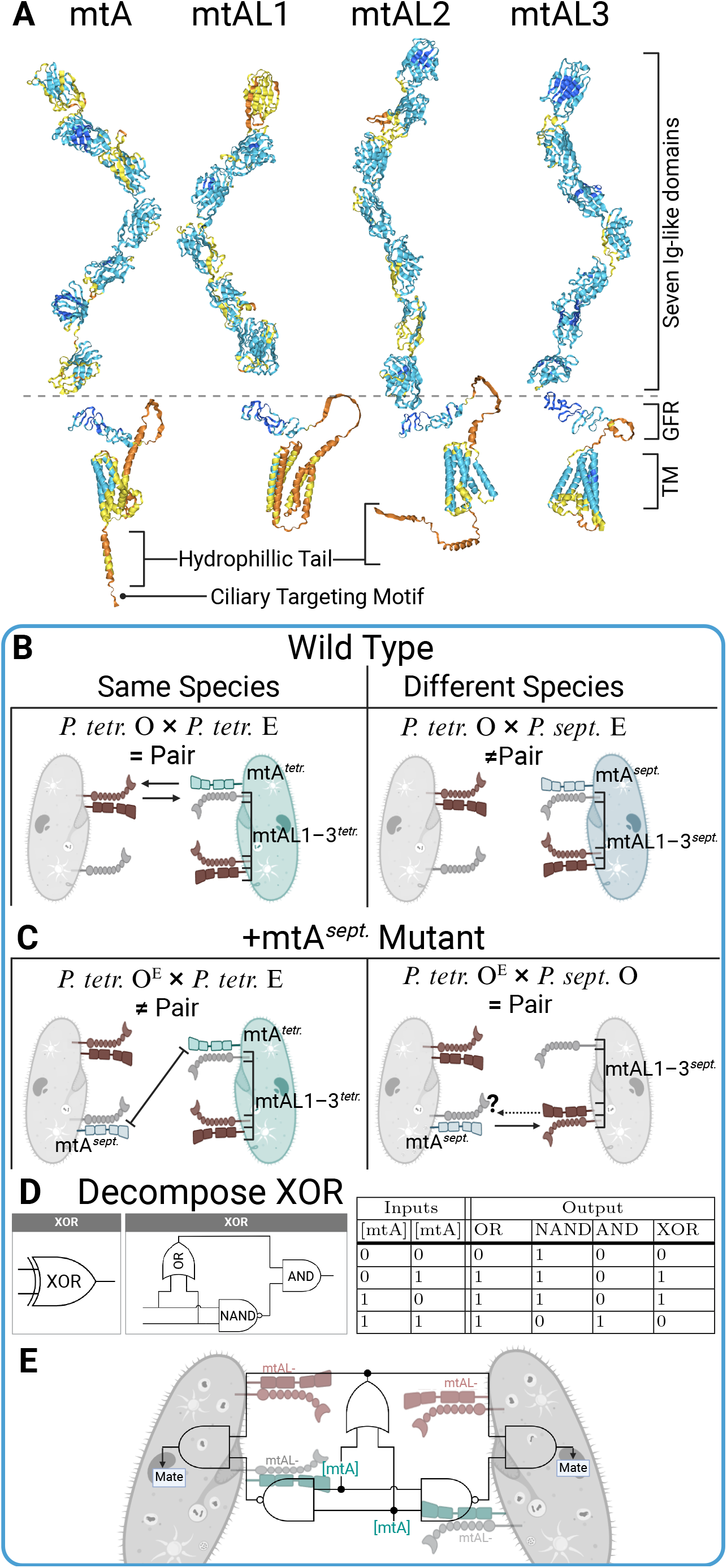
Bidirectional XOR models of *P. aurelia* Even and Odd MT. A) *P. tetraurelia* mtA-Like family sequence-to-structure predictions by AlphaFold3 are generated by partitioned predictions of their mature N-terminal Ig-like domains alongside predictions of their growth factor receptor (GFR) and transmembrane (TM) domains. All labeled structural features appear among predictions of various sequenced *P. aurelia* species. Residues are color coded by pIDDT confidence: High confidence is denoted by pIDDT > 90, (dark blue), confident as 90 > pIDDT > 70 (blue), low confidence as 70 > pIDDT > 50 (yellow), very low confidence as < 50 (orange). B) Interaction model and summary of cross-species agglutination studies between complementary MT of *P. tetraurelia* and *P. septaurelia*. In this model, MT-signaling requires *cis*-dimerization of mtA-Like (brown) receptors that recognize conspecific (left vs. right) mtA in-*trans*. In E type cells, mtA *cis*-dimers with mtA-Like (cyan/blue) may act as MT receptor. C) *P. tetraurelia* O type that are microinjected [38] with the *mtA* gene of *P. septaurelia* are labeled O^E^. They fail to agglutinate with conspecific E type (left) whilst gaining the ability to mate with O type *P. septaurelia* (right). In this model, both mtA of *P. septaurelia* and *P. tetraurelia* inhibit agglutination in-*trans*. D) XOR gate decomposition. E) Agglutination of *Paramecium* is a computation of bidirectional XOR where each cell’s surface expression of mtA is considered as a binary input. Across two cells, mtA binding to its species-specific receptor can actuate OR gating. Dual expression of mtA, of any species, causes homomeric inhibition and actuates NAND gating. In the absence of inhibition, the competitive engagement (AND gate) of mtA to mtA-Like receptors signals compatible MT and agglutination.

Because mtA has cross-species homology, the *P. aurelia* binary MT were historically renamed Even (E) and Odd (O). Inter-species mixing of E and O sometimes results in adhesion and sometimes in conjugation. The reactivity of mtA to multiple receptors supports the concept of receptor-ligand promiscuity in the binary system. In contrast, any *Tetrahymena* MT engage all non-self types with equal preference and do not pair across species, indicating tight exclusionary regulation. In *P*.*aurelia*, interspecies conjugation is highly correlated with adhesion but with discrepancies: interspecies E × O can differ in both adhesion and conjugation from O × E [37]. This indicates that two pairing cells sense two different signals, despite their only difference being a single switch in mtA expression. In agreement with prevailing thought that MT-specificity arose from species-specificity [18], we predict that MT specificity evolved by epigenetic switching of mtA out of an ancient set of unisexual cellular adhesion receptors (Figure 2A).

While organisms can convey binary MT in many ways, the adhesion-like, elongated MT proteins in *P. aurelia* suggest differential intermembrane binding. Classical studies demonstrated that fixed cells can agglutinate with live cells only if they are of complementary MT, consequently activating meiosis as well as homotypic (same MT) pairing in the live cells [36]. While we provide further support for pairing cells to activate via two different receptors, we unfortunately cannot probe mechanisms downstream of pairing in *Paramecium* because signaling is inseparable from adhesion. Pairwise adhesion is intrinsically bidirectional, and signaling appears bidirectional in both MT as well.

An additional issue is that the receptor for mtA must be expressed in both MT, but its identity has not been found. By drawing parallels to *Tetrahymena* MT proteins, it is likely to be among the homologous mtAL family (illustrated in Figure 2B). Two important perturbations of mtA, performed by Singh et al. [38], change the agglutination phenotype. First, the ‘E’ phenotype converts to ‘O’ type when *mtA* is silenced or mutated. Therefore, mtA is self-purifying because there is no way to inactivate self-inhibition, and any mtA that fails to inhibit mating to E type would eventually evolve sterility with O type as well. Second, *P. tetraurelia* and *P. septaurelia* are normally incompatible, but O type *P. tetraurelia* that are microinjected to express *P. septaurelia*’s mtA agglutinate with O type *P. septaurelia* whilst losing the ability to agglutinate with E type *P. tetraurelia* (Figure 2C). This case suggests, at least, that an mtA interaction with its receptor is species-specific.

To better understand how surface receptor concentration computes MT compatibility, we deconstructed the XOR mating phenotype to relate it to simpler protein mechanisms involving two protein inputs on apposing cells. XOR should output a mating signal only if two inputs differ (Figure 1E). It is is self-purifying because it requires both non-self activation and self-inhibition. Mechanistically, protein XOR gates require negative cooperativity of two inputs, either allosteric or built by a dimeric combination of basic receptor-ligand interactions [12]. The most basic gates to construct XOR involves an AND composition of OR and NAND (Figure 2D). Thus the logical construction was realized in *P. aurelia* by in-*trans* binding of mtA to its receptor (OR gating). Then, if OR is able to delineate between species, then NAND must be able to delineate between E/O (Figure 2E). In the *Tetrahymena* system, we propose NAND-like, *trans*-homophillic binding as a mechanism that unlocks the binary system of *Paramecium* to expand to a multiple-MT network.

### Mta/Mtb Dimers Compute NOT EQUALS with Inherited Properties of XOR

We notice that the mating system of *Tetrahymena*, despite different requirements, inherits mechanistic features of XOR from *Paramecium*. For example, the XOR gate of a *P. aurelia* species has the logical property of *self-duality* : swapping the two MT protein inputs has no effect on output, so one cell’s ‘self’ identity is the ‘non-self’ identity from an apposing cell’s perspective. In *Tetrahymena*, this feature is similarly mirrored. However, while *P. aurelia* can delineate self and non-self via a binary switch in mtA expression, *T. thermophila* must distinguish oneself from six forms of non-self (Figure 1D–E). Both cells of *Tetrahymena* must simultaneously present a MT-specific receptor and ligand. Rather than with a single MT-protein, the two proteins represent cellular identity with a dualistic external presenter and internal sensor of self. In contrast to *Paramecium* whose activation cannot be decoupled from adhesion, we exploited the atypism of knockout-imposed asymmetry in MtaΔ ×WT experiments to explore a minimal case where an WT cell’s receptor activates based on two Mtb inputs: one on the same cell and one on its mating partner. Then, self-non-self logic can occur in-*trans* as a result of differences between heteromeric and homomeric affinities.

*Trans*-homodimeric affinity is a common characteristic of cellular adhesion molecules enabled by their large surface area and intermembrane symmetry [26]. We start with the premise that XOR in proteins involves a parallel computation of species-specific OR and MT-specific NAND, and recognize these computations can be satisfied by Mta recognition of a complementary Mtb ligand, and ligand inhibition by *trans*-homomeric Mtb, respectively. To investigate the structure-to-logic relationship between *T. thermophila* MT proteins and computation, we submitted the Mta and Mtb sets of sequences to AlphaFold. In our initial attempts to predict monomeric structures using an earlier AlphaFold2 model [40], we observed that each predicted Mtb—but none of Mta—folded inward to interface with itself (example Figure 3A–B). This is a known in-*silico* artifact in predictions of proteins that are homophilic [41]. Mtb monomers were observed to self-interface their first N-terminal Ig-like domain (D1) to their third (D3), but this interface vanished when either D1, D3, or both were artificially swapped with that from different MT (Extended Data). However, when we submitted the same sequences to AlphaFold3 [34], all folded proteins preferred elongate structures that did not self-interface (Figure 3D). In addition, we recapitulated in AlphaFold3 high confidence *trans*-homodimerization of D1–D3, when two D1–D5 Mtb segments were submitted (Figure 3C). Overall structural comparison of *Tetrahymena* Mta and Mtb families shows the same common features with the *P. aurelia* mtA family (Figure 3D). Also in common with *P. aurelia* mtA, Mtb contains the known [42] ‘VxPx’ sequence in a predicted cytosolic region as a potential ciliary targeting motif. Notably, all *Tetrahymena* Mta proteins also contain a glutamine-proline-rich linker region between D4–D5 which may confer flexibility.

**Figure 3:**
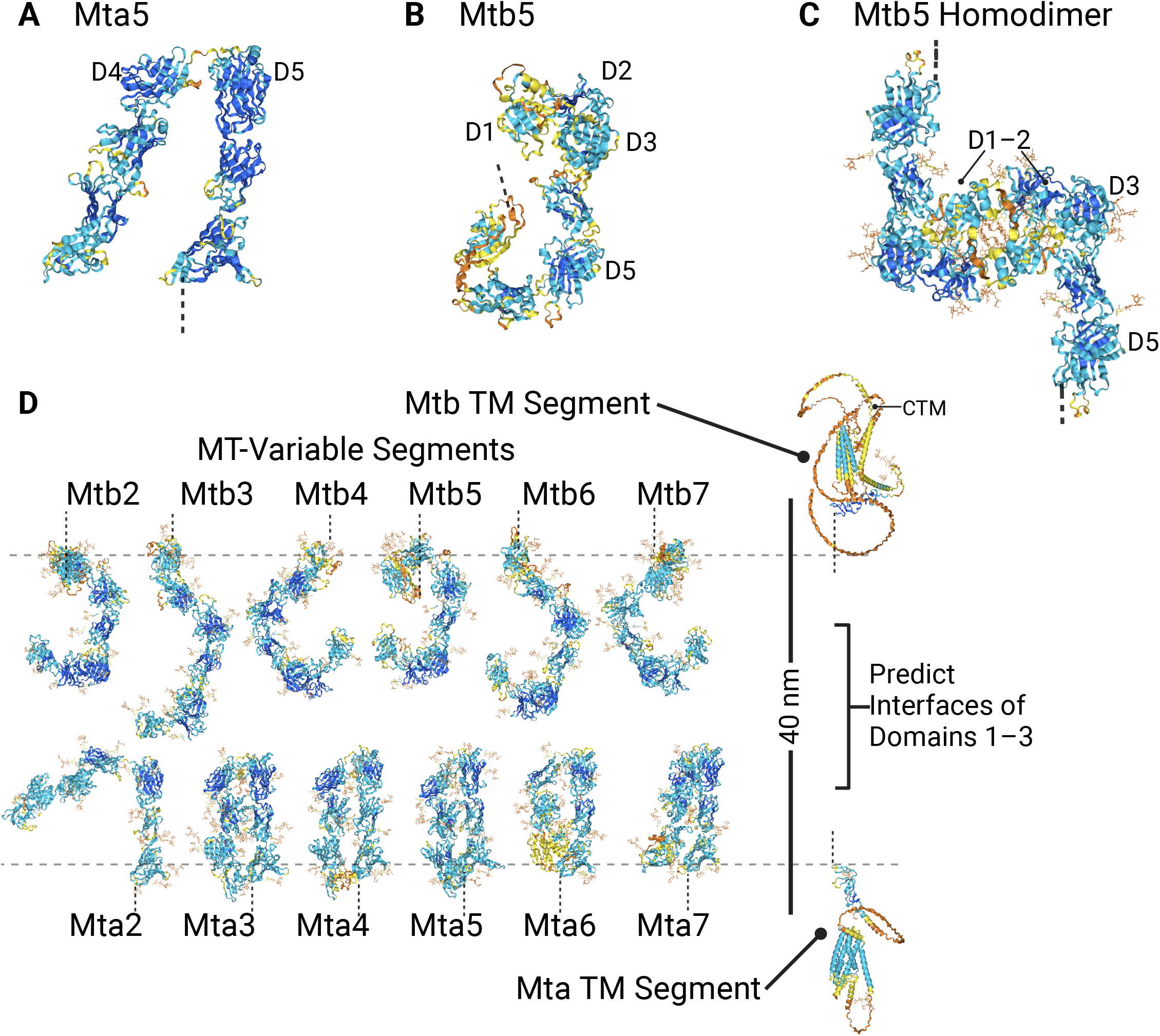
Conserved and homomeric features in predicted structures of *Tetrahymena* Mta and Mtb. A) Example structure of the MT-variable portion of Mta5, all seven β-sandwich domains, generated with AlphaFold2. All Mta proteins are elongate but have moieties divided by a glutamine-proline flexible linker region between the fourth and fifth N-terminal domain (D4–D5). B) Same example procedure for Mtb5. All Mtb proteins are elongate but have D1 self-interfacing with D3. C) Symmetric homodimer prediction of two Mtb5 D1–D5 segments are generated with AlphaFold3 (see Method Details). D) The MT-variable domains of Mta and Mtb sets of structures are predicted by AlphaFold3, drawn to scale with an intermembrane gap distance of 40 nm. Currently in *T. thermophila*, only MTII-VII are sequenced. A potential ‘VxPx’ ciliary targeting motif (CTM) is found on Mtb TM cytoplasmic loops. Residues are color coded by pIDDT confidence: high confidence is denoted by pIDDT > 90, (dark blue), confident as 90 > pIDDT > 70 (blue), low confidence as 70 > pIDDT > 50 (yellow), very low confidence as < 50 (orange).

An important characteristic of MT proteins is their large size and 15-20 nm extracellular length. Adherence of stimulated cells occurs with an intermembrane gap distance of 40 nm [43, 44]. We inferred that the constraints of distance, computational symmetry, and rapid speed of stimulation severely limit the possible solution space of MT computational mechanisms to occur in a single layer—powered by binding kinetics of Mta and Mtb. Because both swimming contact- and adhesion-mediated signaling must be MT-dependent, the structural facets of Mta and Mtb are geometrically consistent with intercellular proteins interfacing near their outer half of N-terminal domains (Figure 3D).

For this reason, we truncated each MT protein to their extracellular D1–D3 domains in the following submission to AlphaFold3 (Method Details). To further test the hypothesis that MT recognition occurs by two MT protein inputs in a receptor-ligand complex (Figure 1B, G), we submitted the truncated sequences of a combinatorial series of Mtb ligands with all Mta/Mtb receptor complexes (Figure 4A). In agreement with Figure 3C, we identified a pattern differentiating homomeric and heteromeric binding: when identical MT are inputted, homomeric interfacing of Mtb-Mtb is competitively preferred over Mtb-Mta (Figure 4A main diagonal). In contrast, some but not all complexes prefer Mtb-Mta heteromeric interfacing when the MT are different (Figure 4A off-diagonals).

**Figure 4:**
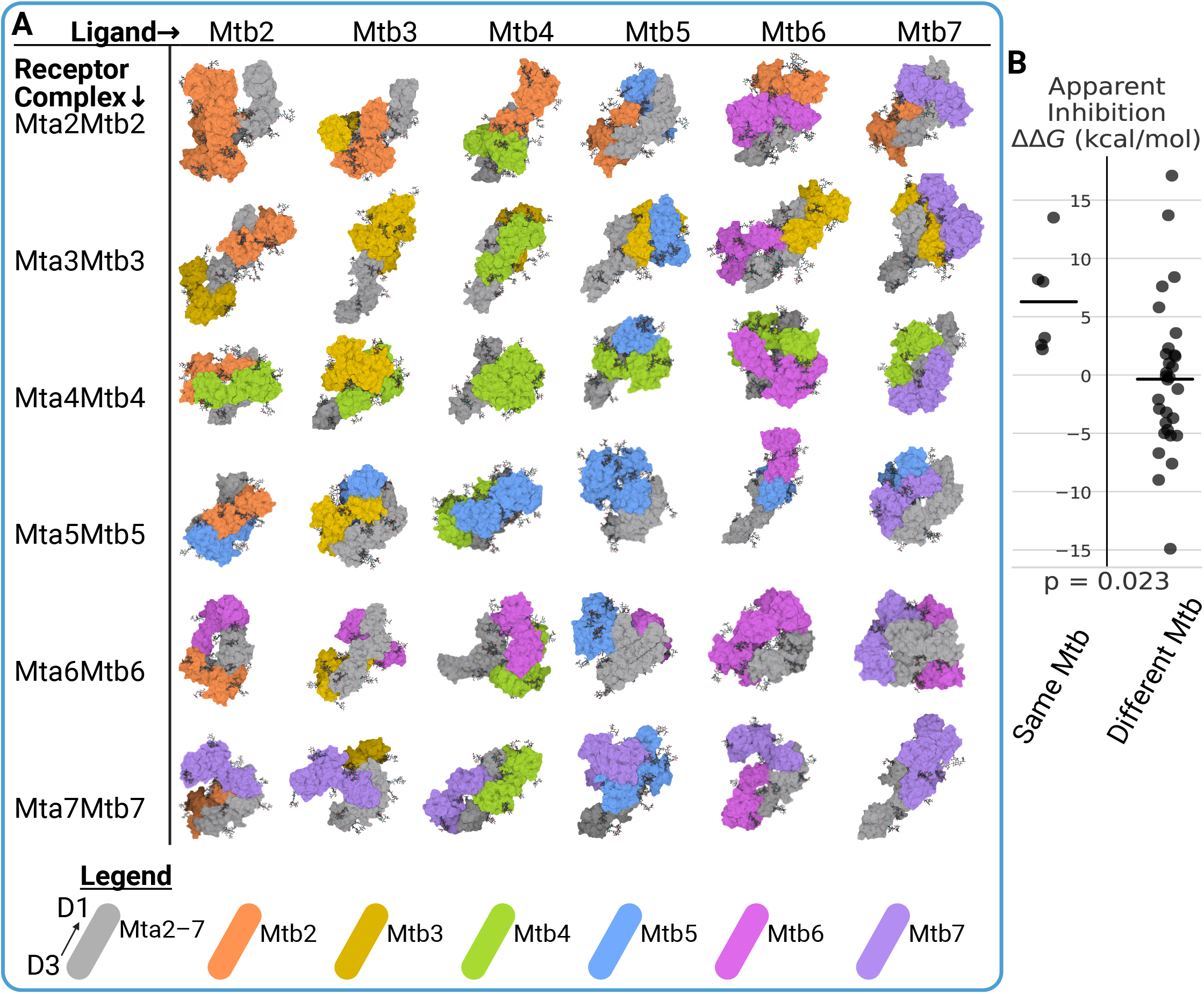
Combinatorial AlphaFold3 predictions of MT complexes all exhibit Mtb-homomeric interfacing and affinity. A) The extracellular first to third domains (D1–D3) from the N-terminus of an Mta/Mtb MT receptor pair (rows), are complexed with a panel of Mtb ligands (columns). Simulation of D1–D3 domains was chosen based on hypotheses of Figure 3. Complexes are all aligned relative to Mta in an upright position from D1 down to D3 (Legend) and visualized using SES representations. N-linked glycans have stick representations. All Mta2–7 are colored gray. Mta2–7 are colored like Fig. 1B. B) Computational prediction of binding energy difference between an Mtb ligand to Mta vs. that to Mtb. ΔΔ*G >* 0 indicate apparent inhibition (Method Details) by Mtb-Mtb interfacing. p = 0.023 for an independence t-test of two samples. N = 6 (Same Mtb) and N = 30 (Different Mtb). Bars indicate means.

To quantify if Mtb homomeric interfacing can inhibit binding to Mta, we compared the contact-based binding energies of Mtb-Mtb vs. Mtb-Mta which is predicted for each complex. All complexes in Figure 4A were submitted to the PRODIGY web server [45], where we recorded the binding energy difference of each ligand to Mta vs. Mtb as a metric of the apparent inhibition of an Mtb inhibitor (Figure 4B). Notably, all cases of identical MT exhibit inhibition averaging 6.3 kcal/mol compared to -0.36 kcal/mol when MT are different (Figure 4B, p = 0.023). AlphaFold-predicted complexes generally suggest a homophilic Mtb ligand to function as its own inhibitor.

Compared to other cell recognition logic, we identified that the ≠ operation features unique properties that agree with empirical observations in *Tetrahymena*. For example, ≠ inherits a key property of XOR in that signaling is broken when either component is knocked out,analogous to the *Paramecium* case with two O-type, mtA^−^inputs. This robust feature is reflected in *Tetrahymena*: knocking out either Mta or Mtb, or both, does not bypass MT-compatibility and thus no cells are allowed to cheat mating by hiding or mutating their receptors. MT protein functionality is intimately tied to their ability to interact correctly with species-specific receptors on partner cells, making them both necessary and sufficient for ≠ logic.

### A Nascent Mating Type Requires Self-confidence and Attractivity

How does a new Mta/Mtb pair expand the preexisting MT system? While *T. thermophila* expresses seven MT, various *Tetrahymena* species have been found to express as little as three, and up to nine MT variants. A comprehensive, mechanistic understanding of MT recognition must also explain the requirements that drive their number and evolution. Although some MT have cross-species homology, some MT appear unique [21]. Also, the absence of a 2^*n*^ pattern of MT variants rules out the possibility that MT computation is a result of a series of binary decisions (as evidenced by the early diverging *P. bursaria* species complex [46]). Overall, phylogenetic evidence suggests that *Tetrahymena* MT evolve recursively. For a system with *n* number of MT, we aim to model the minimal requirements for the *n* + 1^th^ MT.

To model the requirements defining the success or failure of a nascent MT, we demonstrate that binding kinetics between two cells can power a ≠ computation. Because we are interested in probing the differences between self and non-self activation in the pre-stimulation phase, we can assume that two unstimulated cells are phenotypically equal so that only the difference between self and non-self interaction dictates the affinities between receptor and ligand. We assume the implicit assumption of the Hill equation in that simple binding affinity of Mtb to Mta activates an MT receptor (Equation 1). The fraction of a cell’s active receptors, named *P*_active_, is also inhibited in-*trans* by homodimeric Mtb:

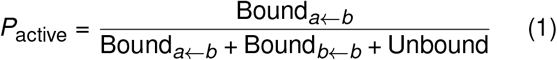

Mta/Mtb are thought to participate as part of an ion-channel signaling complex [33]. Upon contact, we assume mating signal activation depends on the concentrations and method of binding of cell *j* ‘s external ligands to cell *i* ‘s complex. Knowing a monomeric Mtb_*j*_ can function as a ligand, it should activate a MT receptor by binding Mta_*i*_ with a nondimensional equilibrium dissociation constant defined as 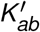. In addition, we make the ideal cooperativity assumption of a symmetric dimer-of-dimers: an MtaMtb complex can engage a juxtaposed partner complex with the summation of both binding energies, 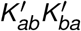 (Method Details). Aforementioned interactions can be inhibited by self-Mtb binding in-*trans* 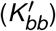. We substitute the nondimensionalized parameters (Equation 2):

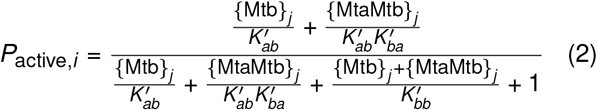

We simplify the signaling model with an assumption of symmetry giving 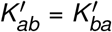, justified when both cells have equal surface concentration of MT receptors and swimming contact confines reactivity to an equal interface of arbitrary surface area. Then we plotted (Figure 5A) the signal response profiles of Equation 2 in a simplified case without monomeric excess Mtb (total surface expression, Mtb_tot_, is adjacent to Mta in-*cis*) and identified that there are in fact two requirements important for MT presentation. The first is species-specific, driven by a low *K*_*ab*_ dissociation constant per all non-self interactions for Mtb to activate other MT. We deem [Mtb_tot_]/*K*_*ab*_ as an ‘attractivity’ ratio. The second requirement is MT-specific, driven by self-inhibition of Mtb by apposing Mtb in-*trans*. Therefore, we deem the [Mtb_tot_]/*K*_*bb*_ as a ‘self-confidence’ ratio. Mta/Mtb affinities underlying the ≠ operation likely incur a trade-off between these two ratios.

**Figure 5:**
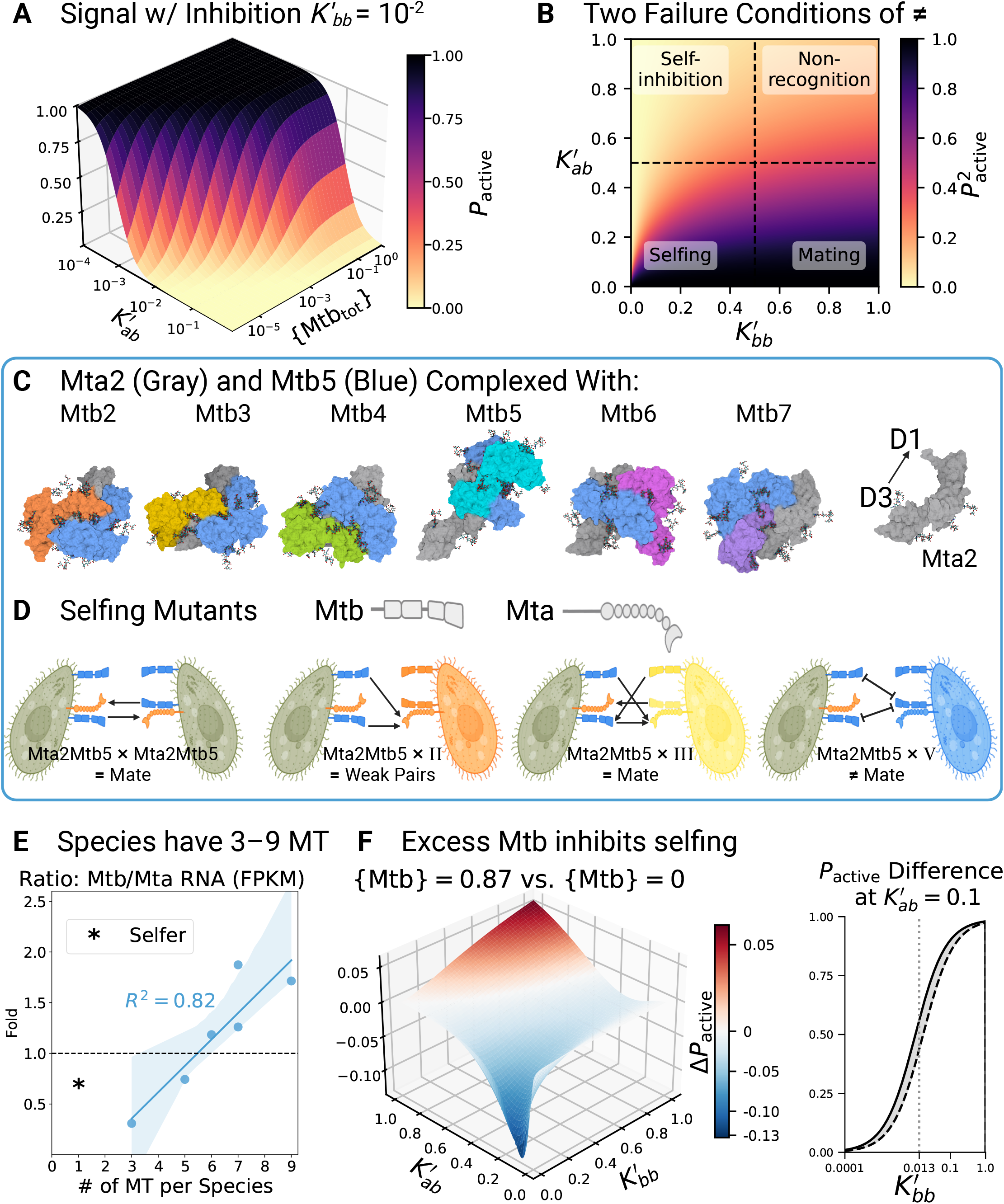
Binding kinetics shows Mta/Mtb chimeric mutants to self-mate whilst failing to react to cognate ligands. A) A lower dissociation *K* value means higher binding energy. MT signaling response curves are activated with strength [Mtb]/*K*_*ba*_ and inhibited by homodimeric *trans*-Mtb with strength [Mtb]/*K*_*bb*_. Equation 2 is plotted when *K*_*bb*_ = 0.01 and in the case of no excess monomeric Mtb so that a dimer-of-dimers of MtaMtb pairs enables positive cooperativity. As two membranes become juxtaposed, 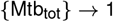. B) Mating requires stimulation of both cells. Contour plot of squared proportions of active receptors 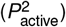. MT signaling requires an inverse relationship between normalized dissociation constants 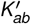 and 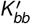. When cells recognize a complementary MT, 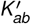 is low and 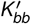 is high allowing mating. When cells are inhibited by homophillic inhibition, 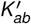 is high and 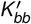 is low. However, selfing occurs when 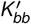 is not sufficiently lower than 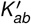 (such as the case of Mta2Mtb5 expressing mutants). C) *T. thermophila* selfing phenotype, caused by the expression of Mta2 and Mtb5, is modeled with the same procedure as Figure 4A. D) Selfing mutant MT of *T. thermophila* that was found by Lin and Yao to express Mta2 and Mtb5 loses the ability to mate with MT-V and MT-II (although loose pairs are reported with MT-II—indicating asymmetrical activation [35]). A mechanistic model in line with Mtb homodimerization competing against Mtb engagement of a complementary Mta is proposed. E) Ratiometric variation across species with *R*^2^ = 0.82 and a confidence interval = 0.95 (shaded blue). According to Yan et al. [21] the mean of multiple values are taken if multiple MT are sequenced. (*) Selfer species (*T. shanghaiensis*) expresses one Mta/Mtb, is considered as ‘unisexual,’ or having one MT, and is not considered. Data is based on starvation-induced RNA-Seq FPKM (Fragments Per Kilobase of exon per Million mapped reads) raw data by Yan et al. [21] F) The overexpression of Mtb as a monomer primarily inhibits selfing. Excess monomeric Mtb was estimated by its ratiometric expression vs. {Mta} in *T. thermophila* according to Figure 5E. Comparing between {Mtb} = 0.87 (dashed line) and Mtb = 0 (solid line), the difference in active receptors from Equation 2 is computed (left). The inhibition (right shaded area) is maximized when 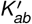 is low and 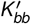 is lower than 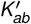 , such that the sensitivity of a MT receptor (right) is maximized around *P*_active_ = 0.5.

**Figure 6:**
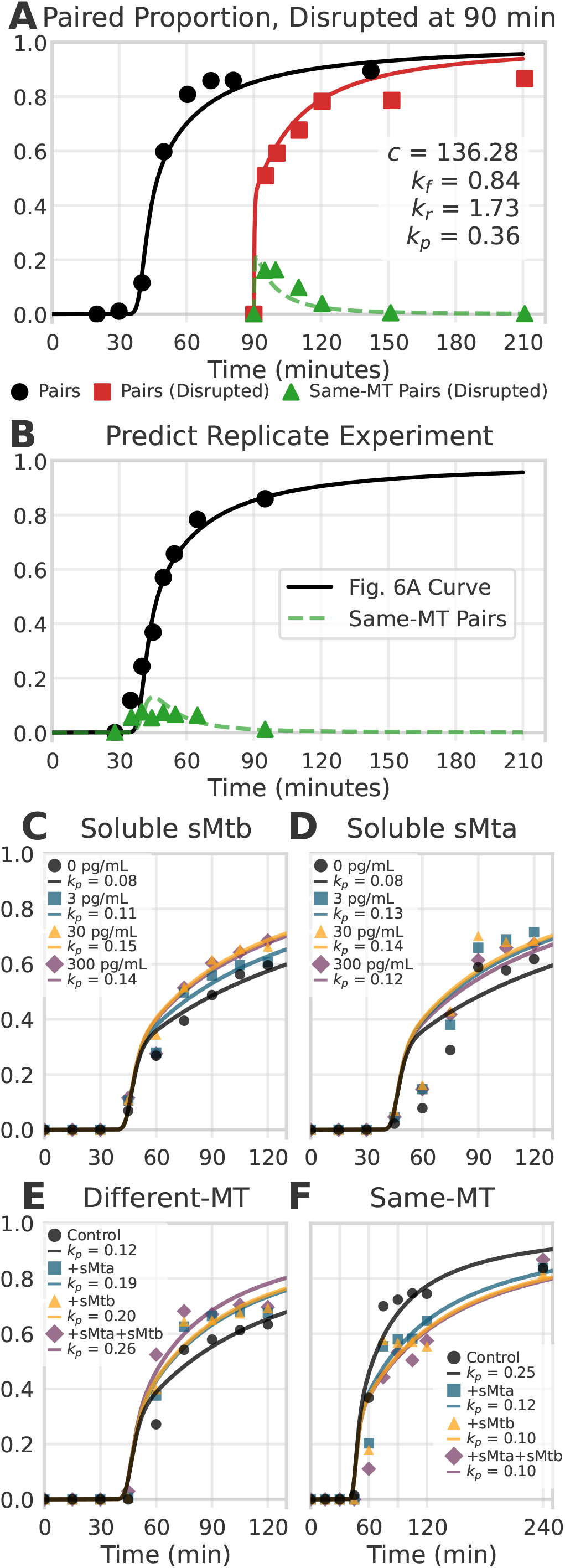
The tractability of pairing dynamics. A) Parametric fitting of required collisions (*c*), reverse pairing rate (*k*_*r*_ ), and MT-compatible catalytic signaling rate (*k*_*p*_), given *k*_*f*_. For Fig. 6A–B, scatter points are digitized from counting experiments by Kitamura et al. [49]. Equal concentrations of starved cells of two complementary MT are mixed. A typical counting (black circle) exhibits a time delay depending on *c* after mixing two MT together. At 90 minutes elapsed when nearly all cells are costimulated, an aliquot is mechanically disrupted by pipette after which cells quickly re-pair (red square). A small proportion of pairs are transient, same-MT pairs (green triangle). B) Simulation of undisrupted cell pairing kinetics using identical parameters from Figure 6A. Proportions of same-MT pairs are counted (green triangle) along with total pairs (black circle). Note that pairing scatter points are typically underestimates of pairing efficiency by a factor related to cell stoichiometric counting precision and cell viability. C) The parameter values *c, k*_*f*_ , and *k*_*r*_ obtained from Figure 6A, are co-plotted with averages of counting experiments (for scatter points Fig. 6C–F) as described by Yan et al. [33] with modifications to *k*_*coll*_ as a result of differences in cell concentration and cell size (Method Details). The mating signal catalytic rate, *k*_*p*_ , was optimized for kinetic pairing among cells pre-incubated with purified soluble (s) extracellular portions of Mtb of complementary MT. D) In the same procedure as Figure 6C, parameter values from Figure 6A are plotted with variable *k*_*p*_ , with poorer fit, to soluble extracellular complementary Mta. E) Combinations of preincubation with 30 pg/mL sMta or 30 pg/mL sMtb, or both, increase *k*_*p*_ additively when soluble protein is complementary MT. F) When 30 pg/mL sMta or 30 pg/mL sMtb (or both) is of the same MT, preincubation decreases *k*_*p*_.

**Figure 7:**
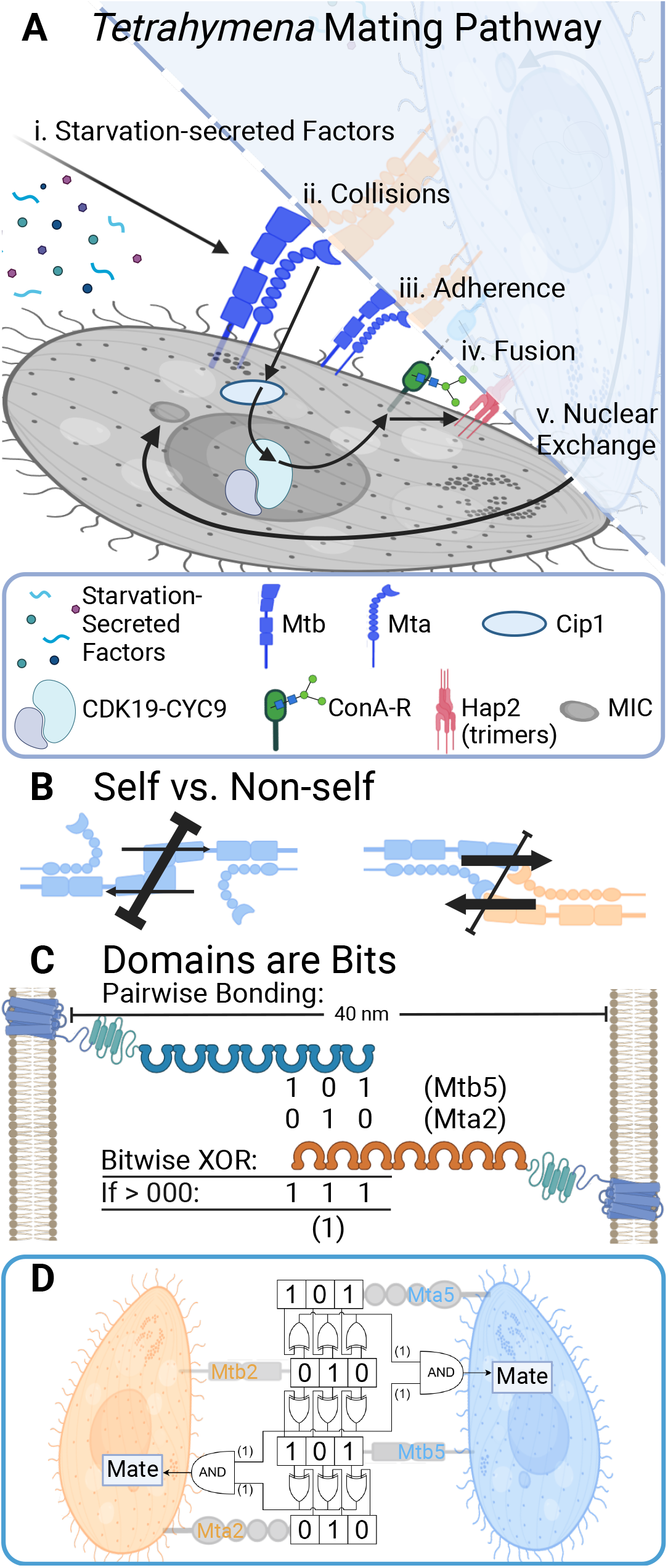
Synthetic biology prospects of *Tetrahymena*. A) Overview of mating pathway schedule and duration: (i. 4 hr) Starvation induces the secretion of MT-nonspecific signal factors [22]. (ii. 0.5–1.0 hr) MT-specific collisions, buffered by Cip1 regulatory subunit of CDK19-CYC19 (Cyclin-Dependent Kinase-CYClin) [47], induce Mtb/Mta/ConA-R/Hap2 expression and morphological tipping. (iii.) Positive feedback is likely mediated by pairing. (iv. ∼1hr) Membrane fusion requires bilateral expression and function of Hap2 as well as other costimulatory factors [56]. (v. ∼5–6 hr) Formation of the conjugation junction while the micronucleus (MIC) performs meiosis, then pronuclear exchange and pair separation occurs. B) Dimer-of-dimers cartoon of ≠ kinetic responses to depend on the activation both receptors and the strength of inhibition. C) Binary encoding of MT protein interactions shows, at minimum, 3 bits of extracellular information transmission. Assuming at least a decade difference in 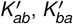 , and 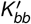 can dictate self vs. non-self, pairwise bonding manifests a bitwise XOR operation. Note only a 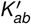 interaction of decimal 2 and 5 is drawn as an example. D) Dimer-of-dimers of Figure 7C. A mating type receptor complex requires two non-self inputs (AND gate).

To quantify the two requirements that lead to mating following adherence (when {Mtb_tot_} = 1), we plotted the trade-off of 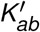 vs. 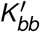 (Figure 5B). The objective of a ≠ operation is to mate with all ‘non-self’ identities and to inhibit only self-mating, but insufficiently low 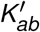 can cause general failure in intercellular recognition. More importantly, when 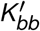 is insufficiently lower than 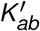 , then a MT cannot inhibit inbreeding occurring when pairs form of the same-MT. This observed phenomena in *Tetrahymena* is known as “selfing,” and is found in a small percentage of natural isolates and cultured cells in that they are able to mate with their own clones.

Although there are several inducible mechanisms of the selfing phenomena [47], selfing failure in cells of the same-MT can be directly linked to MT proteins. For example, the rare species *T. shanghaiensis* expresses only one MT that apparently fails to inhibit selfing. Moreover, Lin and Yao [35] describe mutant cell lines that express a MT chimera of Mta2 and Mtb5 to exhibit selfing. To predict the interactions of this chimeric MT, we submitted the sequences of Mta2/Mtb5 complexed with a panel of ligands to AlphaFold3 with the same procedure as in Figure 4A. Similarly to complexes in Figure 4A, Mtb5 exhibits homomeric interfacing with Mtb5 (Figure 5C). The pairing assays by Lin and Yao [35] describe the selfing cells expressing Mta2/Mtb5 to also have lost the ability to pair with MT-V. Additionally when Mta2/Mtb5 are mixed with MT-II, only weak pairing is observed in a manner that phenocopies “MtaΔ × WT” pairings. We modeled these findings according to our homomeric inhibition mechanism (Figure 5D). If we were to assume that Mtb binds to Mta by only pairwise receptor-ligand interactions, then the Mta2Mtb5 chimera response to either MT-V or MT-II should be interchangeable—but that is contrary to what is observed.

The curious observation of asymmetrical stimulation in MtaΔ knockouts demonstrates that presentation of Mtb, without Mta, can sufficiently function as a ligand. This feature may be inherited from a *P. aurelia*-like binary system, but also may point towards MT-expansion. Various *Tetrahymena* species have been found to express 3–9 MT with no phylogenetic relationship dictating the number of MT (sequenced by Yan et al. [21]). Interestingly, we plotted the ratio of RNA-Seq Mtb/Mta expression and found a positive trend (*R*^2^ = 0.82) in the ratiometric excess expression of Mtb with the number of MT per species (Figure 5E). Theoretically, each *MTA/MTB* gene pair is able to modulate their expressions because their upstream sequences are also MT-specific. MT genes are in a linear arrangement and all but a single pair are excised before MT presentation. An *MTA/MTB* gene pair is oriented head-to-head so that their transcription is closely coupled. Moreover, otherwise similar architecture in their genetic location, coordinated low expression, protein structure, and membrane targeting allow us to assume that excess RNA is positively mapped to excess surface protein expression.

We hypothesized that higher Mtb presentation provides a nascent MT a selective advantage by increasing both ‘self-confidence’ against self-mating and increasing ‘attractivity’ to established MT. We compared activation (Equation 2) with excess ligand ({Mtb} = 0.87 according to the relative expression in *T. thermophila*), to zero excess when MT interactions occur only in a dimer of dimers (as in previous Figure 5A–B). According to Equation 2, excess Mtb primarily provides a selective advantage against selfing (Figure 5F at low 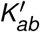 and low 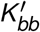 ) with the greatest inhibition when 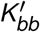 is lower than 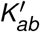. Therefore, monomeric Mtb would maximize discrimination between self and non-self by inhibiting formation of self dimers-of-dimers. Although not modeled by Equation 2, if Mtb monomers are also able to inhibit apposing Mtb, there would likely be some diminishing returns of excess ligands—representing a type of substrate inhibition that may constrain the maximum number of MT.

Beyond the selfing conundrum, decomposing a ≠ gate into a dimer-of-dimers may also explain a conundrum observed in dual-MT expressing mutants. Lin and Yao [35] describe a mutant that expresses both MT-IV and MT-VI simultaneously and is able to mate with all MT, including itself. However, Yan et al. [33] reported a transformed MT-VI cell line with an exogenous overexpressed Mtb2-eGFP that is able to mate with itself but loses the ability to mate with MT-II. The latter pairing points to overexpressed Mtb2-eGFP having an inhibitory function competitive against Mta6/Mtb6 presentation. We find that *Tetrahymena* discriminates necessarily with affinity, benefited by ratiometric expression.

*Tetrahymena* selfing can occur by self-inhibition failure, a mechanism we argue to be driven by MT-specific affinities. To compare our characterizations of MT-systems, we find that *P. aurelia* differs in that it conveys binary MT with variation only of surface concentration. While *P. aurelia* may be constrained to binary MT, they are benefited in that they are not found to ever self-mate with the same MT. Here, we have described the protein mechanics underlying MT computation that reconciles several lines of empirical evidence. What is missing is a ‘chimera-of-chimeras’ pairing (e.g. generating Mta2/Mtb5 × Mta5/Mtb2), with data which would fall under the non-recognition quadrant of Figure 5B and could triangulate the differential affinities of 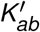 and 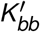 between self and non-self. Overall, we have assumed that Mta-Mtb interact in a one to one molar ratio, and that binding kinetics on both sides functionally constitutes a dimer-of-dimers of MT proteins. To justify several assumptions made of MT interactions, we continue to model pairing and signaling as a dynamical system.

### Reaction Kinetics of Mating Estimates its Signal Speed

We reaffirm that mating is not a simple process; it requires tight coordination of two complicated biological cells. However, we aim to extract and model only the physical and MT-specific aspects of the mating process to verify assumptions about Mtb/Mta mechanics. The largest difference between *Paramecium* and *Tetrahymena* mating is that *Paramecium* cluster/agglutinate immediately when complementary MT are mixed together, while *Tetrahymena* only pair after 30-60 minutes of physical collisions between complementary cells. The process, named costimulation, is observed on the anterior tips of swimming cells with surface expression of an adhesive glycoprotein, named Concanavalin A Receptor (ConA-R), as well as morphological rounding. Hence, while *Paramecium* MT signaling cannot be easily decoupled from adhesion, *Tetrahymena* MT computation occurs in a separate pre- and post-adherence phase.

When two MT are mixed together, they swim in 3D space and collide randomly. The pairing process can be analogized to bimolecular reaction kinetics but, as found by varying concentrations/proportions of mixed cells [48], the elapsed time required for pairing does not correlate with cell concentration but rather with the amount of expected complementary collisions. This correlation is consistent with MT signaling and adhesion expression to require multiple transient physical contacts. For instance, a concentration of 500,000 cells/mL swimming at a speed of 450 µm/sec (∼10 body lengths per second) and with a width (collision diameter) of 15 µm will expect an average 101 collisions/hr to a complementary MT. However, only a fraction of collisions results in pairing via their anterior tips, which we model with a proportionality factor comprising their adhesive surface area.

We assign the former parameter as collision frequency *k*_coll_, and the latter pairing rate as a forward reaction rate *k*_*f*_ , which is reversible by *k*_*r*_. In addition, we denote MT-positive signaling as *k*_*p*_ , which represents MT-specific catalytic conversion from adherent to MT-compatible pairs and thus eventual commitment to mating. Herein, we separate MT-signaling parameters *k*_coll_ and *k*_*p*_ from adhesion parameters *k*_*f*_ and *k*_*r*_. We then simulate mass action kinetics to model cell and pair concentrations in a dynamical system with *X* and *Y* single cells of different MT as well as *M*_2_ committed pairs (Equation 3):

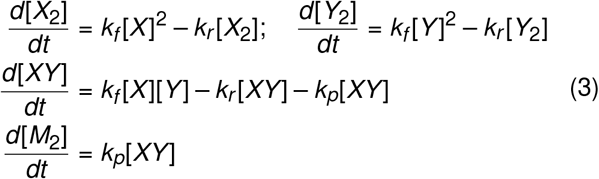

Here, *X*_2_ and *Y*_2_ represent weakly adhered cells of the same MT, while *XY* are weakly adhered cells of different MT. We apply the simplification *X* = *Y* , since usually equal proportions of cells are mixed together. We estimated *k*_*coll*_ and *k*_*f*_ from collision theory (Method Details). To incorporate a 30–60 minute delay in costimulation, we assume that *X* and *Y* cells do not react until they reach stimulation by a constant threshold (*c*) number of collisions, which are random events accumulating over time (Method Details).

We fitted the remaining model parameters *k*_*r*_ , *k*_*p*_ , and *c* to classical, microscopic counting experiments by Kitamura et al. [49] who capture initial stimulation time and total pairs as well as rapid repairing after pair disruption by pipette (Figure 6A). Crucially, they also distinguish and count pairs of the same-MT (Figure 6A-B), which are weak and transient.

Our model correctly predicts the slower formation of such pairs in undisrupted cells (Figure 6B). We also model an average 136 collisions to be necessary for stimulation, as well as a mating commitment catalytic rate, *k*_*p*_ , of 0.36 min_-1_. We show that MT-signaling can occur across single cells with repeated contacts which last seconds, and among adhered pairs that last a few minutes.

To test the signaling effect of MT protein concentration on the *M*_2_ catalytic rate *k*_*p*_ , we digitized plots recently published by Yan et al. [33] and applied the kinetic parameters optimized previously (*c* = 136, *k*_*r*_ = 1.73 min_-1_). To match their experimental conditions, *k*_*f*_ was calculated with different collision parameters: 20 µm width estimate from images vs. 15 µm from that of Kitamura et al. [49], and *X* = *Y* concentrations at 250,000 vs. 500,000 cells/mL, respectively. Importantly, Yan et al. [33] record pairing of cells that were preincubated with recombinant, soluble, extracellular portions of Mta and Mtb. When cells were preincubated with soluble Mtb (sMtb), we observed a saturable, concentration-dependent relationship of sMtb with *k*_*p*_ (Figure 6C). A similar relationship is exhibited by sMta although the modeled parameters from Figure 6A correlated poorly with all sMta kinetic experiments (Figure 6D). At equal concentrations, sMta and sMtb had similar effects on *k*_*p*_ and when incubated with both, an additive effect was observed (Figure 6E-F).

Although we cannot explain the molecular details of how sMta and sMtb interact with membranes, their equal effects on *k*_*p*_ suggest that Mta and Mtb signal at a 1:1 molar ratio. Yan et al. finds that sMta/sMtb does not stimulate expression of the ConA-R adhesion receptor [33], suggesting additional membrane/cytoskeletal components are necessary for signaling. Our model aligns with only *k*_*p*_ being affected, maintaining the possibility that sMta and sMtb stabilize (or disrupt when same-MT) a ‘dimer-of-dimers’ signaling complex that naturally occurs within a 40 nm gap distance. Interestingly, *k*_*p*_ of control experiments (Figure 6C-D, 0 pg/mL) is calculated to be near half the saturation levels of *k*_*p*_ (Figure 6C-D, 300 pg/mL). If we were to assume a Monod relation between the *kp* phenotypic parameter, adhesion surface area [50], and the activation of MT-receptors, then 0 pg/mL of soluble Mtb would correspond to a wild type affinity ratio [Mtb]/*K*_*ba*_ ≈ 1, meaning surface concentrations already maximize the sensitivity of our derived Hill curves (Figure 5A) and thus the sensitivity delineating non-self from self. However, further experiments with direct measurement of paired active and inactive MT complexes are necessary to validate our inference.

## DISCUSSION

### Limitations of this Study

We have approached MT non-self and self molecular recognition with an inductive series of cases, progressing from a binary-MT base case in *P. aurelia* to the variable natural number of MT in *Tetrahymena*, and to the system requirements and stoichiometric methods for MT expansion. Although we have recapitulated findings with in-*silico* modeling and deconstructed the logical underpinnings of Mtb and Mta that reconcile all previous studies, this study does not explain the atomic principles that would provide the structure and function of an Mtb/Mta receptor. The combinatorial complexity, low abundance, and large size of paired-MT complexes would pose challenges for structural studies.

Knowing that four MT-specific proteins are involved in a *Tetrahymena* pair, we have derived a demonstrative XOR model for *Paramecium* (Figure 2D). However, the difficulty of binary XOR comes from the multitude of multilayered mechanisms that possibly mediate cell pairing, especially considering that the identity of the receptor of mtA is unknown. Since the receptor’s expression is MT-non-specific, it could be deduced that in E-type cells, mtA inhibits its receptor in-*cis*. A similar deduction can be made that *Tetrahymena* Mta necessarily inhibits Mtb in*cis*. Both deductions were discussed by previous studies on *P. aurelia* [38] and *T. thermophila* [35]. If *cis*-inhibition were MT-specific, it would also explain selfing failure in mutant *T. thermophila* expressing Mta2/Mtb5. Outside of findings that align with our results of *trans*-homophillic binding of Mtb (such as overexpression of Mtb2-eGFP inhibiting MT-VI compatibility with MT-II), our primary criticism of this hypothesis in *Tetrahymena* is that signaling would require additional steps that do not provide an apparent advantage over simple pairwise receptor-ligand affinity.

### One-to Two-to Many-Mating Type Systems

In fact, an engineered receptor-ligand system would only require pairwise affinity to non-self MT plus aversion to self MT, which calls into question what would drive it to evolve the more complicated interactions that we have described. *Trans*-homophillic binding likely confers robustness to a sexual system that is *generatively designed*, rather than engineered. From an evolutionary perspective, a nascent MT confers a 1/*n* fold selective advantage in mating, where *n* is the preexisting number of MT. Then the evolutionarily stable strategy is to encode random expression of the maximum number of intercompatible MT, beholden to MT-protein physical limits in affinity themselves. A nascent MT must not mate with itself as that could manifest a system collapse back into a unisexual species.

As the “physical substrate” of sex, MT proteins are understandably constrained in their structural evolution and diversification, yet their genetic expression is much more plastic. For example, various *P. aurelia* species exhibit at least five different regulatory mechanisms to switch mtA_+_ vs. mtA_–_ [37] (but curiously never of the mtA receptor). Compared to single switching, we have demonstrated how the MT-specific switching of a receptor-ligand pair provides the increased complexity in interactions that beget multiple MT. Mta and Mtb are the only recognized homologs of each other among *Tetrahymena* pointing towards duplication and diversification, although their precise evolution is still in question. Meanwhile, mtA and mtaL are the homologs among *Paramecium*, and so it remains unknown if MT-specificity by Mtb/Mta arose either independently or consequentially from mtA. Hypothetically, heterologous dual replacement of Mtb/Mta (or rather their extracellular domains) with combinations of *Paramecium* mtA/mtAL1-3 could reconstitute a binary system in *Tetrahymena*. If tested to be true, it would identify the *Paramecium* receptor for mtA (Figure 2B), cement a dimer-of-dimers architecture, and may evidence either *trans*-inhibition or *cis* depending on the result of co-expressed combinations.

Mtb/Mta are predicted to be elongate, heavily glycosylated proteins with a large surface area on β-sandwich domains to enable molecular recognition. Functionally speaking, Mta, Mtb, and mtA are unnecessarily large for their purpose compared to MT-pheromones, which can be small molecules or peptides. In contrast to pheromonal, binary yeast mating which exhibits concentration- and population-dependent behavior [51], collision-mediated signaling in *Tetrahymena* is independent of population ratios of MT. The difference is further exemplified in multiple-MT systems that require non-self pheromones to over-power autocrine self pheromones [25]. In a similar light to how the yeast mating pathway has elegantly aided our understanding of cell decision-making via the integration of noisy signals [52, 53], the *Tetrahymena* system offers future insights into affinity-driven, self-non-self signal robustness among single cells.

### Barcoding Promiscuity vs. Exclusivity

The Mtb/Mta family is unlike those comprising multi-cellular growth factors (and their analog to MT pheromones) that operate as promiscuous receptor-ligand networks. Mtb/Mta operates with mutually exclusive expression and interaction specificity in a manner that bridges the conserved features of development with the random variation of antigens. Understanding these mating antigens and their underlying circuitry can enhance our study of *Tetrahymena* mating behavior (Figure 7A). For example, the symmetric expression of both an MT-specific receptor alongside a ligand may inform several perplexing aspects of *Tetrahymena* sex. Reciprocal signaling may allow the decoupling of MT-signaling from non-specific adhesion. Similar to the coordinated cell wall breakdown in yeast [54]), *Tetrahymena* evolved a mechanism of co-commitment prior to fusion that prevents irreversible mixing of cellular identity and maintains their viability for future mating partners. Co-commitment is exemplified by the symmetric requirement of ConA-R for adhesion as well as the tightly regulated bilateral functional requirement of Hap2, a virus-like membrane fusogen typically expressed haphazardly and asymmetrically when only in sperm [55]. In addition, several recently identified costimulatory factors are necessary for *Tetrahymena* pairing and fusion [56]. Although the identity for ConA-R (and its cognate) is yet unknown, its cytoskeletal coupling [57] may mediate morphological tip transformation. Lastly, knockout of the recently discovered, *Tetrahymena*-specific inhibitory adaptor Cip1 causes selfing by bypassing MT-signaling [47]. We have summarized *Tetrahymena* mating with central relevance towards MT-recognition (Figure 7A). Ongoing inquiry in the field will continue to illuminate its pathway mechanisms.

A dimer-of-dimers architecture (Figure 7B) requires 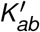 and 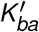 only relatively lower than 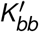 for six non-self interactions and vice versa exclusively for self interaction. Totalling seven MT, *T. thermophila* must transmit cellular identity across 40 nm with a minimum of 3 bits of information. The multi-bit configurability is definitely afforded by a transition from concentration-to affinity-driven signaling. Herein, we propose a theory of self vs. non-self by a binary representation of MT decimal number that is encoded along Mtb/Mta domains (Figure 7C). Pairwise bonding between Mtb to Mta and Mtb to Mtb are then physically programmed as bitwise XOR where each bit defines, say, at least a decade difference in *K*_*′*_. Then affinity can be protein-engineered with as simple as a series of hydrogen bonding interactions [4]. One feature of this representation is ‘backwards compatibility’ as a new MT would simply increase the bit width (e.g. MT-VIII = 1000) of the system.

It has been said that, “Nothing in immunology makes sense except in the context of infectious diseases and self-nonself discrimination” [58]. How does the relatively domestic Mtb/Mta set inform larger design challenges in synthetic biology and immunoengineering? For instance, *trans*-homodimers of ICAM-1 (InterCellular Adhesion Molecule-1) agglutinates circulating tumor cells and aids in their metastatic adhesion and migration [59]. Also, the engagement of integrin receptors of ICAM-1 stimulates neutrophils as an effective innate immunotherapy [60]. There is some underlying logic: neutrophil regulate *trans*-adhesion to ICAM-1 by their expression of inhibitory ICAM-1 in-*cis* [61]. Thus neutrophils express a ligand alongside its receptor, perhaps to modulate signaling with concentration-dependence or prevent neutrophil self-adhesion. As a general mechanism, the coexpression of any *trans*-homophilic ligand could inhibit activation of a receptor complex.

XOR logic may also inform mechanisms pathogens use to expand their variant antigenic repertoires and also aid us in the design of vaccines to be both immunogenic and multivalent. Both antigen and MT protein have functional and discriminatory requirements to activate (non-self) vs. inhibit (self) immune/mating activation, while still having an advantage in varying their epitopes. Like variant pathogenic antigens, the mutually exclusive expression and linearly repeated genetic structure of *MTA/MTB* pairs make them conducive to duplication and evolution: any two Mta and/or Mtb exhibit structural conservation despite peptide-scale variability (averaging ∼60% sequence similarity) [18]. Similarly, strains of the malaria parasite *Plasmodium falciparum* evolved 45–90 *var* genes which encode large, structurally-conserved adhesion molecules. These antigens exhibit modular variation with recombination of domain cassettes to accomplish cell type adhesion specificity with combinatorial dual-binding to ICAM-1 and EPCR, or ICAM-1 and CD36, or they bind and agglutinate other erythrocytes. The unique expression of EPCR + ICAM-1 binding domains contributes to its specific inflammatory adhesion to activated neutrophils and to human brain endothelial cells, the latter recognized as the primary determinant of malaria mortality [62].

In conclusion, we reveal how simple protein interactions can give rise to multi-bit cellular decision making, thus expanding our understanding of the evolutionary limits of protein computation. We also propose several experimentally testable mechanistic predictions that take advantage of the elegant logic of mating. To demonstrate the biologically feasible design requirements of XOR and ≠ on protein interfaces, future applied work could duplicate and engineer the *n* + 1^th^ type. Our findings bridge the gap between Mtb/Mta expression and cellular behavior, providing an explanation for long-observed phenomena in *Tetrahymena* mating.

## Acknowledgements

This paper commemorates the retirements this year of Paul O’Reilly, Donna M. Cassidy-Hanley, and Theodore G. Clark, who had provided invaluable mentorship to R.A. We thank Matthias Garten for inspiring discussions, feedback, and generous access to Biorender.com. Figures were created with Biorender.com. We thank members of the Garten (Parasite Biophysics) Lab for feedback. We further thank feedback and comments on the manuscript from Donna Cassidy-Hanley and Theodore Clark. This material is based upon work supported by the National Science Foundation Graduate Research Fellowship Program to R.A. under Grant No. DGE-2146755. Any opinions, findings, and conclusions or recommendations expressed in this material are those of the authors and do not necessarily reflect the views of the National Science Foundation. We also acknowledge the support of R.A. by a Sang Samuel Wang Fellowship (part of Stanford Graduate Fellowships in Science and Engineering), and a Nagel (Tau Beta Pi) Fellowship. We further acknowledge the financial support for M.P. (CZI BioHub, HHMI Faculty Fellowship, Schmidt Foundation, Moore foundation, NSF CCC Grant).

## Author Contributions

R.A. conceived, designed and performed the experiments and analyzed the model/simulations. M.P. conceptualized, validated, and supervised the work. R.A. wrote the software and the original draft manuscript. M.P. and R.A. reviewed and edited the manuscript.

## Declaration of Interests

The authors declare no competing interests.

## STAR METHODS

### Key Resources Table

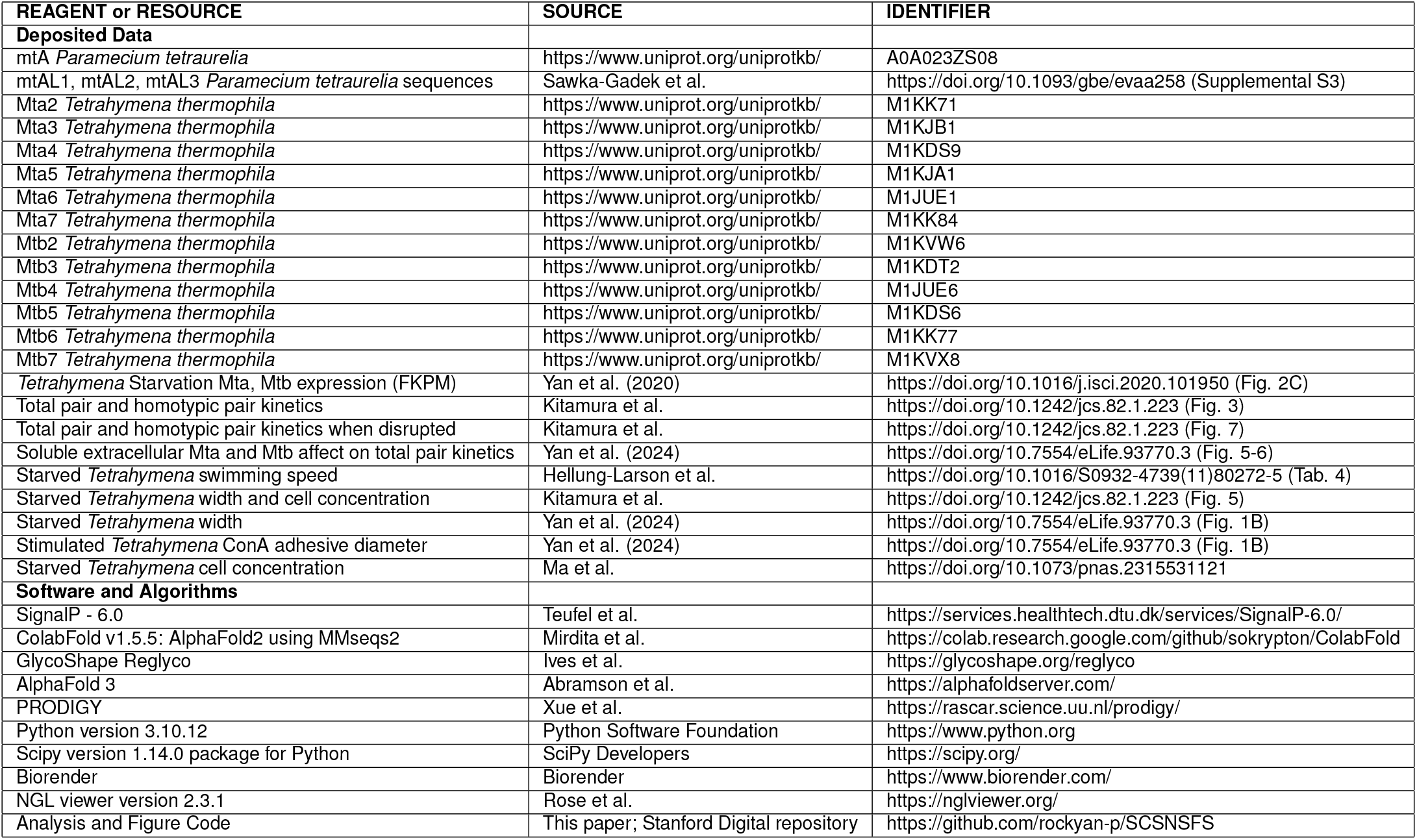

### Resource Availability

#### Lead contact

Further information and requests should be directed to and will be fulfilled by the lead contact, Rocky An.

#### Materials availability

This study did not generate new unique reagents.

#### Data and code availability

All original code, including data used to generate all quantitative figures, will be deposited in the Stanford Research Data collection in the Stanford Digital Repository and will be publicly available upon the date of publication. For ease of access, code is also available on GitHub at https://github.com/rockyan-p/SCSNSFS. Code is categorized by its associated figure. As of the latest date of creation, the entire study can be replicated and reanalyzed using the deposited code and web-server input files. (Extended Data): contains confidence metrics and those structures of AlphaFold2 monomeric predictions mentioned.

### Method Details

#### Nomenclature

Due to the interdisciplinary nature of the study, a mixture of nomenclatures are used which are to be explained in this section. The capitalization for *Paramecium* proteins (mtA) differ from *Tetrahymena* (Mta). *T. thermophila* mating types are named in Roman numerals (e.g. MT-II). Before sequencing methods could identify *Tetrahymena* and *Paramecium* species, sexually isolated MT-intercompatible groups were defined a species syngen as part of a species complex. The *P. aurelia* complex and *T. pyriformis* complex are the relevant historical examples.

#### Structural and Complex Prediction Pipeline

In the study of antigen-antibody interactions as well as the protein engineering field, binding energy is the discriminant connecting protein structure to function. We employed AlphaFold3—which excels at atomistic accuracy and antibody-antigen interactions in particular [34]—with the hypothesis that it incorporates these intrinsic general features of adhesion proteins in predictions of Mta/Mtb interfaces.

Protein sequences for *P. tetraurelia* were obtained from Sawka-Gadek et al. [37], and for *T. thermophila* from Cervantes et al. [19]. Only MT-II through MT-VII Mta/Mtb have been sequenced. Mature protein sequences were obtained by cleaving predicted N-terminal signal peptides using SignalP-6.0 [63]. For monomeric AlphaFold2 predictions, sequences were submitted to AlphaFold2 using MMseqs2 [40] on Google Colab, without templates. For monomeric AlphaFold3 predictions, mature sequences were initially submitted to AlphaFold3 [34], then each sequence is partitioned at its seventh globular domain and the two partitions folded seperately. Glycosylation is essential for cell pairing, so *T. thermophila* Mta/Mtb N-linked glycosylation sites were predicted by submitting the initially-folded iterate to GlycoShape Re-glyco [64]. *Tetrahymena* N-linked glycans are relatively homogeneous and non-complex: the majority of N-glycans are high-mannose in the glycoform of Man3-GlcNAc2 to Man5-GlcNAc2 [65]. For final predictions, sequences were repopulated with Man5-GlcNAc2 to predicted sites and resubmitted to AlphaFold3.

For complex multimer predictions, Mta/Mtb full sequence lengths sums over the maximum limit of AlphaFold3, thus combinations of truncated D1-D3 domains were submitted. To ensure reproducibility in the comparison of folded complexes, the random seed of input files was set to 2, 3, 4, 5, 6, or 7 depending on the MT number of the corresponding mtA. NGL viewer [66] was used for cartoon visualizations with confidence metric-colored B-factors. Biorender.com was used for SES (solvent-excluded surface) 3D representations.

#### Binding Energy Predictions Pipeline

Binding energies of Mta to a cognate Mtb ligand vs. Mtb to a cognate Mtb ligand were compared in all 36 folded complexes. Each interface energy was calculated using the PRODIGY (PROtein binDIng enerGY prediction) web server [45]. In the trimeric complex, is it assumed that competitive binding of Mtb to Mtb vs. Mtb to Mta can inhibit signaling, thus the difference in their binding energies is calculated as a metric of inhibition. Quantification and comparison between groups was performed (Quantification and Statistical Analysis). Likely contributing to variation in the data are the predicted confidence metrics of the MT complexes being generally lower than that of monomers (Extended Data), with the Mtb5-Mta5-Mtb5 complex having the highest confidence of note. If—as in a few cases with Mta3—there are no predicted contacts between two protein chains, binding energy was estimated to be zero. In the same-MT cases, there are two identical Mtb and thus two possible interfaces to Mta, thus the one with most affinity is used as a conservative measure of its inhibition. Binding energy raw data is available in the Figure 4B code file.

#### Kinetic Derivation

Although the kinetics powering self-non-self signaling is derived from only one cell’s perspective and then symmetry is applied, it still necessitates careful consideration of the interaction’s directionality at the interface. Therefore, cell *j* is denoted the ligand-presenter and cell *i* the receptor-sensor. Because this receptor is known to contain Mta and Mtb in-*cis*, Mta-Mtb is denoted as a dimeric receptor (D) in either unbound (D), bound-activated (A) or bound-inactivated (I) states. The equilibrium dissociation constants are defined by the method of Mtb ligand binding to the dimeric receptor. Also, an assumption of dilute reactants allows nondimensionalization of concentrations to “moles of receptor per cell” reference units:

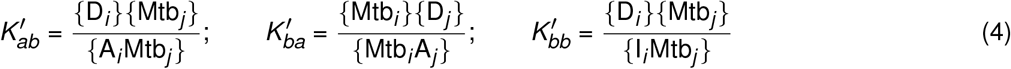

The Hill equation measures the activation response as the fraction of bound (active) states out of overall states. It implicitly assumes activation by binding without considering intermediate states or activation energies, and would not change the functional form of Equation 2 if such are considered. In this case, there are two possible activatory states, two inhibitory states, and one unbound state. Either/both Mtb_*j*_ , as a monomer, or D_*j*_ as a dimer, can function as a ligand or inhibitor.

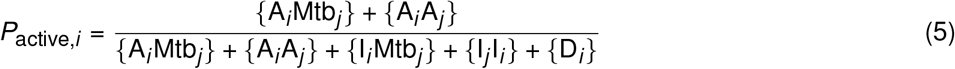

To estimate the affinity of dimer-of-dimers interactions for the ‘A_*i*_ A_*j*_ ‘ state, an ideal proximity-induced cooperativity is assumed by summing two dimeric binding energies for the symmetric complex. The ‘I_*i*_ I_*i*_ ‘ state still only exists by single binding between Mtb_*i*_ and Mtb_*j*_.

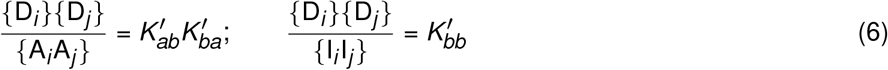

Finally, Equation 2 is obtained by substitution and factoring {*D*_*i*_ }:

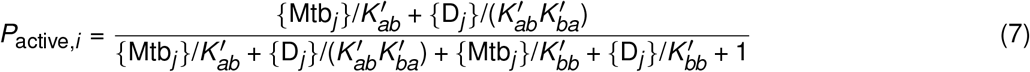

In all Figure 5 plots *K*_*ab*_ = *K*_*ba*_ was assumed, and per-cell nondimensional concentrations are to approach [Mtb_tot_] = 1 as the membranes between two cells become juxtaposed. Mating strictly requires stimulation of both cells [67] so Figure 5B assumes mating strength as a function of the product of 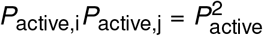 with i, j symmetry.

#### Collision and Costimulation Derivation

The swimming cell collision frequency (*k*_coll_) on a per-cell basis was estimated by analogy to molecular collision theory, accounting for cell area, concentration, and relative swimming speeds. A steric factor gets *k*_*f*_ from *k*_coll_:

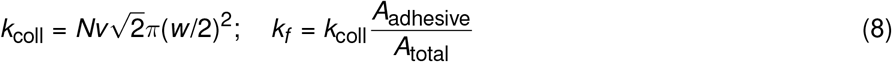

where *N* is the cell concentration of the complementary MT, *v* is the swimming speed, and *w* is the cell width. Cell size, count and speed can depend on growth and starvation conditions in culture. The cell width was estimated as 15 µm or 20 µm from figure images by Kitamura et al. [49] and Yan et al. [33], respectively. Cells are also seen to adhere when ConA-reactive tips collide. The reactive area was estimated by images of the pairing synapse and fluorescent ConA [33]. The steric factor is then the adhesive tip of circular area with diameter 15 µm, over a total surface area 4*π*(*w* /2)_2_.

According to experimental conditions of Kitamura et al. [49] and Yan et al. [33], *N* is 5 × 10^5^ cells/mL and 2.5 × 10^5^ cells/mL respectively, which are high densities that saturate chemotaxis. Starved cells are approximated to swim at 450 µm/sec [68]. Under starvation conditions, *T. thermophila* transition from run-and-tumble like motility of average length ∼1 mm to a straighter swimming motility of length ∼10 mm [29, 32]. Befitting the molecular collision model, with a *k*_coll_-predicted mean free path length of 8 and 9 mm respectively, cells are thus assumed to swim comparably straight.

After mixing cells, some time (*t* ) is required for cells to costimulate. The number of mean collisions *λ*_*t*_ advances at a rate *dλ*_*t*_ /*dt* = *k*_coll_ {*X*} , where *X* is the proportion of non-paired cells. Paired cells do not significantly collide, an assumption well-justified according to McCoy [48], because paired cells lose velocity and sink. At early times, collision count is a Poisson random process, which when *λ*_*t*_ is large approximates a Normal distribution with the same mean, and variance *λ*_*t*_.

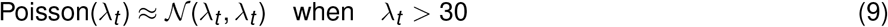

The normality is maintained even as *dλ*_*t*_ /*dt* decreases. It is assumed that a fixed threshold number of collisions *c* stimulates a cell. Out of all cells, {*S*} is defined as the stimulated proportion of cells that can participate in the pairing reactions. Then S is calculated from the cumulative distribution function of the Normal distribution:

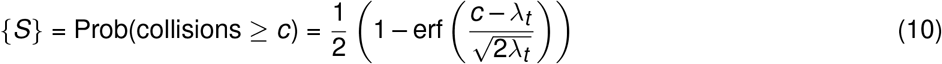

where erf() is the error function.

### Quantification and Statistical Analysis

For Figure 5E, *MTA* and *MTB* expression data of *Tetrahymena* species that have multiple-MT were obtained from averages reported by Yan et al. [21]. The fold difference in expression was calculated from the reported FPKM (Fragments Per Kilobase of exon per Million mapped reads) values, and plotted as a linear regression with shaded 95% confidence interval. The selfer species *T. shanghaiensis*, although expression plotted, was not included in the analysis because the cells of same MT mate among themselves. In the analysis in Figure 5F, the excess expression of *T. thermophila* in Figure 5E was used with the assumption of similar post-transcriptional context between Mtb and Mta so that excess expression approximately correlates with excess monomeric Mtb. Qualitatively speaking, higher relative expression of *MTB* is consistent across multiple *T. thermophila* sequencing efforts [67].

#### Parameter Optimization

Figure 6 scatter data was manually digitized from published figures (Key Resources Table). Mass action kinetics based on Equation 3 and Equation S7 were simulated with the Scipy.odeint module. The Scipy.optimize module was used to perform a least squares optimization strategy to get parameters *k*_*r*_ , *k*_*p*_ , and *c* to fit all scatter series in Figure 6A. However, the nonlinear optimization does not guarantee the best solution and is sensitive to initial condition estimates (in our case as 1 min^−1^, 1 min^−1^, 100 respectively). Nevertheless, differences in solutions are minor so that they do not affect any outcomes about conclusions made from *k*_*r*_ , *k*_*p*_ , and *c*. Because *k*_*r*_ and *c* are defined in per-cell units, they are treated as constants among *T. thermophila* and their optimized values from Figure 6A are applied to Figure 6B–F. Thus they are assumed independent of the different experimental conditions across the plots. For example, while stimulation may be reversible because of ConA-R and MT-receptor turnover, the timescale of all experiments is similar where virtually all cells stimulate within ∼1 hour.

#### Binding Energy

For each of 36 complexes of MtaMtb2-7 with Mtb2-7, competitive inhibition was hypothesized to occur by competition of the Mtb binding interface. The strength of inhibition is approximated by the relative interaction energy: ΔΔ*G* = Δ*G*_*a*←*b*_ – Δ*G*_*b*←*b*_ , where Δ*G* is the negative-valued free energy change for each interface. To confirm significance (p < 0.05) in inhibition differences between same-MT (N = 6) and different-MT (N = 30) groups, an independent two-sample t-test was performed. The assumption of normality of the groups was confirmed using the Shapiro-Wilk test. Both tests were performed using the Scipy.stats module.

## Notes

https://github.com/rockyan-p/SCSNSFS

### Competing Interest Statement

The authors have declared no competing interest.

